# Intrinsically Dominant Conformational Diversity in PDZ1 within the Tandem PDZ1-PDZ2 of Human Syntenin-1 Underlined by Crystal Structures

**DOI:** 10.1101/2025.05.22.655525

**Authors:** Natsuno Ando, Yuya Hanazono, Koya Sakuma, Nobutaka Numoto, Takeshi Tenno, Atsunori Oshima, Nobutoshi Ito, Hidekazu Hiroaki

## Abstract

The intrinsic dynamic asymmetry between homologous PDZ domains in multidomain scaffold proteins offers critical insights into evolutionary mechanisms enabling multivalent partner recognition. Through systematic X-ray crystallographic analysis of human syntenin-1’s PDZ1-PDZ2 tandem, we resolve nine high-resolution structures that uncover fundamental differences in conformational plasticity between these sequentially similar domains. Pairwise root-mean-squared deviation (RMSD) analysis of 20 PDZ1 structures across multiple crystal forms reveals substantial structural variability concentrated in the Lys119-Ile125 and Ala181-Glu184 loops - key regions governing ligand specificity within PDZ1’s binding cleft. In stark contrast, PDZ2 maintains remarkable structural conservation across all crystallographic environments, indicating divergent evolutionary constraints on these tandem domains. Crucially, comparative analysis of isotropic B-factors demonstrates their inadequacy in capturing the full scope of conformational heterogeneity, emphasizing the necessity of multi-structure comparisons for mapping dynamic landscapes.

Molecular dynamics (MD) simulations implemented through GROMACS corroborate these crystallographic observations, showing elevated residue-specific fluctuation (RMSF) values in PDZ1’s ligand-binding interface compared to analogous PDZ2 regions. This consistency across experimental and computational approaches confirms that PDZ1’s conformational diversity represents an inherent biophysical property rather than crystallographic artifact. The observed dynamic asymmetry suggests a functional division of labor: PDZ1’s structural plasticity enables broad ligand recognition via conformational selection mechanisms, while PDZ2’s rigid architecture likely stabilizes the tandem domain arrangement. These findings provide an atomic-level rationale for syntenin-1’s pleiotropic roles in cellular signaling and establish a structural blueprint for developing domain-selective therapeutics. Given syntenin-1’s clinical relevance in cancer metastasis, viral pathogenesis, and neurodevelopmental disorders, our work advances strategies for selectively modulating PDZ1-mediated interactions while preserving PDZ2’s scaffolding functions through structure-guided inhibitor design.

**Highlights:**

◊ Crystal structures of human Syntenin-1’s PDZ1-PDZ2 tandem reveal intrinsic conformational plasticity in PDZ1, particularly in ligand-binding loops, contrasting with PDZ2’s rigid architecture
◊ Pairwise RMSD analysis of 20 PDZ1 structures demonstrates substantial structural variability in the Lys119-Ile125 and Ala181-Glu184 loops, key regions governing ligand specificity
◊ Molecular dynamics simulations confirm that PDZ1’s conformational diversity is an inherent biophysical property, not a crystallographic artifact
◊ The asymmetric dynamics between PDZ1 and PDZ2 suggest a functional division: PDZ1’s plasticity enables broad ligand recognition while PDZ2 stabilizes the tandem arrangement
◊ These findings provide a structural basis for developing domain-selective Syntenin-1 inhibitors with potential applications in cancer metastasis, viral pathogenesis, and neurodevelopmental disorders

## Introduction

Syntenin-1 (SDCBP/MDA-9) is a 32 kDa cytosolic scaffold protein that serves as either an adaptor or a critical hub in cellular signaling and trafficking networks [Grootjans, J. J., et al., 1997]. The architecture of syntenin-1 features two (Postsynaptic density-95/Discs large/Zonula occludens-1 (PDZ) domains in tandem (PDZ1 and PDZ2) flanked by N- and C-terminal regions, enabling it to act as a versatile protein-protein interaction hub. While syntenin-1 is extensively studied in context of its role in exosome biogenesis through interactions with syndecans and ALIX [Baietti, M. F., et al., 2012], emerging evidence highlights its other broader functions, including synaptic regulation [Hirbec, H., et al., 2005], cytoskeletal dynamics [Zimmermann, P., et al., 2001], receptor trafficking [Zimmermann, P., et al., 2005], and viral entry [Gordon-Alonso, M., et al., 2012]. These activities are linked to its divergent subcellular localization, which spreads from the plasma membrane, focal adhesions, endosomal compartments, to the early secretory pathway.

A key feature of syntenin-1 is its ability to coordinate signaling complexes via PDZ domain-mediated interactions. The PDZ tandem binds diverse partners, including Frizzled receptors [Egea-Jimenez A. L., et al., 2016], phosphatidylinositol 4,5-bisphosphate (PIP2) [Zimmermann, P., et al., 2002], and viral proteins [Jimenez-Guardeño, J. M., et al., 2014], through cooperative mechanisms that enhance binding avidity. For example, PDZ2 engages in tripartite interactions with the Frizzled 7 C-terminus and PIP2, leveraging adjacent binding pockets to stabilize membrane-associated signaling platforms [Egea-Jimenez, A. L., et al., 2016]. Such interactions underpin syntenin-1’s role in non-canonical Wnt signaling and syndecan recycling, processes critical for cell polarity and growth factor responses [Luyten, T., et al., 2008].

Involvement of syntenin-1 in many important biological processes highlights syntenin-1 as a target for drug discovery. Thus, for example, atomic-resolution structures of syntenin-1’s PDZ domains are critical for rational drug design. The X-ray crystal structure of PDZ1 bound to the inhibitor KSL-128018 (PDB code: 6AK2) reveals a noncanonical binding mode involving residues beyond the canonical groove, including Trp-4 and Chg-2, which engage in hydrogen bonds and hydrophobic interactions [Kang, B. S., et al., 2003]. Similarly, PDZ2’s PIP2-binding pocket, adjacent to the peptide groove, is stabilized by water-mediated networks [Zimmermann, P., et al., 2002]. These structural details may explain how syntenin-1 achieves high-affinity interactions with diverse partners and highlight targetable regions for disrupting oncogenic signaling.

Accordingly, the extended binding interfaces and hydration networks between PDZ1 and PDZ2 position syntenin-1 as a promising therapeutic target. Inhibitors such as KSL-128114, designed to leverage these structural features, demonstrate nanomolar affinity, metabolic stability, and efficacy in glioblastoma models [Haugaard-Kedström et al., 2021]. Targeting syntenin-1’s PDZ-mediated interactions with syndecans, Frizzled receptors, or viral proteins (e.g., SARS-CoV-2 E protein) could mitigate cancer metastasis, neurodevelopmental disorders, and viral pathogenesis, respectively [Boukerche, H., et al., 2005][Kegelman, T. P., et al., 2014] [Gordon-Alonso, M., et al., 2012]. Thus, high-resolution structural data of individual PDZ domains may further enable the development of domain-selective inhibitors to modulate syntenin-1’s multifunctional roles in disease.

Initially, we aimed to resolve the drug-PDZ1 complex structure for anthranilic acid derivatives, analogs of Dishevelled PDZ inhibitors [Hori, K. et al,, 2018]. While co-crystallization attempts with these compounds were unsuccessful, we determined ten high-resolution apo-form crystal structures of the syntenin-1 PDZ1-PDZ2 tandem including one crystal of PDZ1-PDZ2 tandem complexed with compound E5 (PDZ2i) [Hoffer, L. et al., 2023]. Structural analysis revealed significant conformational variation in PDZ1’s canonical ligand-binding pocket, contrasting with PDZ2’s static architecture. Here, we evaluate conformational divergence across 20 PDZ1 structures from the ten independent crystals and compare them to PDZ2. GROMACS-based molecular dynamics (MD) simulations of isolated PDZ1 and PDZ2 domains further highlighted intrinsic flexibility in PDZ1’s Lys119-Ile125 and Ala181-Glu184 loops.

## Materials and Methods

### 1. Expression and Purification of Protein Samples

The DNA fragment encoding the human syntenin-1 PDZ tandem (residues 113-273 and 113-276, see Figure 1) was amplified via polymerase chain reaction (PCR) and subsequently cloned into a glutathione-S-transferase (GST)-fusion expression vector, which included an PreScission™ Protease (Cytiva, Tokyo, Japan) cleavage site. Protein expression was induced by the addition of 1 mM isopropyl β-D-1-thiogalactopyranoside (IPTG) in *Escherichia coli* BL21-CodonPlus (DE3)-RIL Competent Cells ((Agilent Technologies Japan, Ltd., Tokyo, Japan). The PDZ tandem was expressed in LB medium, and cell pellets were subsequently lysed by sonication. The lysate was clarified through centrifugation and purified by affinity chromatography using COSMOSIL® GST Accept (nacalai tesque, Inc., Kyoto, Japan) in a buffer containing 50 mM Tris-HCl (pH 7.5), 150 mM NaCl, and 1 mM EDTA. The GST-tagged protein was then subjected to cleavage using PreScission™ Protease at 4℃. After complete digestion, the cleaved PDZ tandem was separated from GST and further purified by gel filtration chromatography using HiLoad™ 26/600 Superdex™ 75 prep grade (Cytiva). The protein-containing elution was subsequently dialyzed in a buffer composed of 25 mM HEPES (pH 7.4), 150 mM NaCl, and 1 mM DTT.

**Figure 1:**
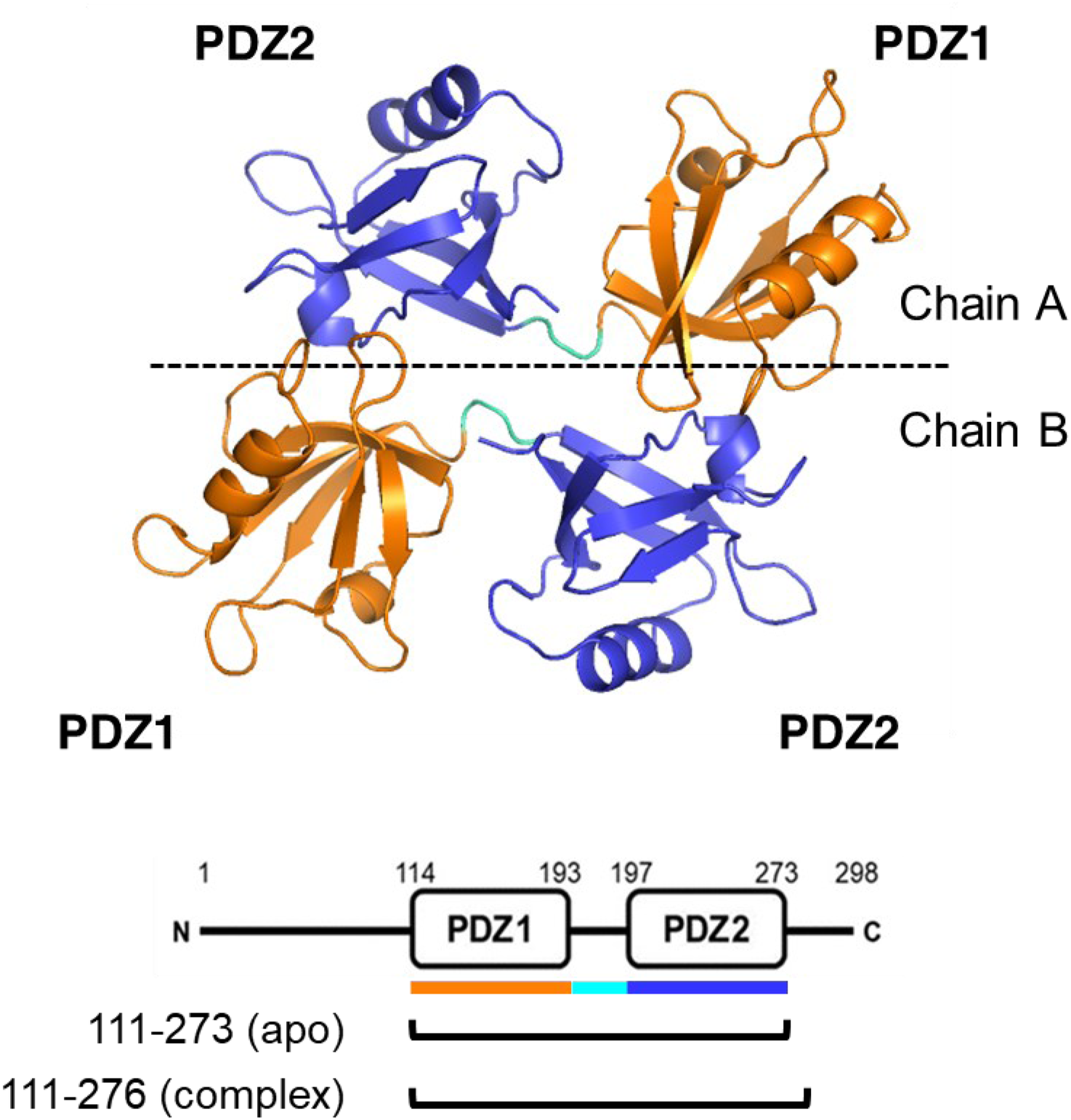
Crystal structure and domain structure of the apo-syntenin-1. The crystal structure contains two molecules of the syntenin-1. The PDZ1 domain is assigned Glu114 to Arg193 (orange), and the PDZ2 domain is assigned Arg197 to Phe273 (blue). 163 amino acids construct was used to determine the structure of apo-syntenin-1. 166 amino acids construct was used to determine the structure of syntenin-1-compound complex.

### 2. Crystallization and Data collection

Crystallization experiments were conducted using the sitting drop vapor diffusion method at 20℃. The specific crystallization conditions for the apo-syntenin-1 PDZ protein are detailed below: In this study, we initially intended to obtain the crystals of the complex between the PDZ tandem and the anthranilic PDZ inhibitors, NPL1010, 3005, 3026, and 3027. The chemical structures of the compounds are summarized in Supplementary Figure 1.

- **S1 (PDB code: 9VA6)**: Crystals were obtained from a solution containing 5 mM NPL3005, 0.2 M Sodium chloride, 0.1 M HEPES pH 7.5, and 25% w/v Polyethylene glycol (PEG) 3,350.
- **S2 (PDB code: 9VA9)**: Crystals were obtained from a solution containing 5 mM NPL3005, 0.2 M Sodium formate, and 20% w/v PEG 3,350.
- **S3 (PDB code: 9VAC)**: Crystals were obtained from a solution containing 5 mM NPL3026, 0.2 M Ammonium acetate, 0.1 M Sodium citrate tribasic dihydrate pH 5.6, and 30% w/v PEG 4,000.
- **S4 (PDB code: 9VAD)**: Crystals were obtained from a solution containing 5 mM NPL3027, 0.2 M Lithium sulfate monohydrate, 0.1 M Tris pH 8.5, and 25% w/v PEG 3,350.
- **S5 (PDB code: 9VAF)**: Crystals were obtained from a solution containing 5 mM PDZ1i, 0.2 M Sodium chloride, 0.1 M BIS-TRIS pH 5.5, and 25% w/v PEG 3,350.
- **S6 (PDB code: 9VAI)**: Crystals were obtained from a solution containing 0.2 M Ammonium acetate, 0.1 M Sodium acetate trihydrate pH 4.6, and 30% w/v PEG 4,000.
- **S7 (PDB code: 9VAL)**: Crystals were obtained from a solution containing 0.2 M Sodium chloride, 0.1 M Tris pH 8.5, and 25% w/v PEG 3,350.
- **S8 (PDB code: 9VB9)**: Crystals were obtained from a solution containing 5 mM NPL1010, 0.1 M BIS-TRIS pH 5.5, and 25% w/v PEG 3,350.
- **S9 (PDB code: 9VBB)**: Crystals were obtained from a solution containing 5 mM NPL1010, 0.2 M Ammonium acetate, 0.1 M BIS-TRIS pH 5.5, and 25% w/v PEG 3,350. The crystallization condition for the syntenin-1 PDZ protein-compound complex is outlined below:
- **S10 (PDB code: 9V90)**: Co-crystallization was performed in a solution containing 0.5 mM PDZ2i, 0.4 M ammonium acetate, 0.1 M Sodium acetate pH 4.6, and 20% PEG 3,350.

All crystals were grown from a protein concentration of 6 mg/mL. Crystals were cryo-protected by immersion in mother liquor supplemented with 10% (v/v) glycerol before freezing. Diffraction data for syntenin-1 PDZ crystals were collected at the Photon Factory BL-17A and processed using X-ray Detector Software (XDS), [Kabsch., W. et al, 2010]. The phase problem was solved by molecular replacement using the structure of PDB code: 8BLU as a template with the program PHASER [McCoy, A. J. et al., 2007]. Models were refined using Phenix [Adams, P. D. et al., 2010]. The figures were prepared using PyMOL software (http://www.pymol.org/).

### 3. Molecular Dynamics

#### Preparation of Initial Structures

The PDZ1 and PDZ2 domains were defined as residues 113-193 and 197-273, respectively, and extracted from the original crystallographic structure to serve as initial coordinates for MD simulations. To ensure consistent construction of simulation boxes for both PDZ1 and PDZ2 domains, the PDZ2 domain was structurally aligned to the PDZ1 domain using the “align” command in PyMOL, and this superposed model was subsequently used as the initial structure. GROMACS (version: 2024.5) [Van Der Spoel, D. et al., 2005] was employed as the MD engine, utilizing the charmm36-jul2022.ff force field, a GROMACS adaptation of charmm36 (Huang, J. et al., 2017). Throughout all simulation procedures, hydrogen atoms were constrained to their ideal geometry using the riding model, and a time step size of 2 fs was applied.

#### Simulation System Construction

The initial protein structure was embedded within a dodecahedron simulation box. Prior to solvation, the system was energy-minimized in vacuum using the steepest descent algorithm for a maximum of 500,000 iterations to eliminate any steric clashes between atoms. The minimized model was then solvated with the TIP3P water model, whose geometry was provided as spc216. To achieve a neutral net charge, the system was ionized with 0.1 M-equivalent NaCl-derived ions, and the solvated system underwent a second round of energy minimization. Finally, the system was subjected to a very short NVT simulation (1 ps at 300 K) with harmonic restraints applied to all heavy atoms to ensure the absence of any severe residual steric clashes.

#### Equilibration and Production Run

A 2-fs time step was consistently used in all MD simulations. The system was equilibrated through sequential 100-ps NVT and subsequent NPT simulations at 300 K and 1 bar, with heavy-atom restraints for each trajectory. Different random seeds were used to generate initial velocities for each independent run. Five independent 200-ns production runs were performed for each of the isolated PDZ1 or PDZ2 domains. The final 160 ns of each trajectory was subsequently extracted for detailed analysis.

#### Fluctuation Analysis

The root-mean-squared fluctuation (RMSF) was computed for each Cα atom from frames sampled every 100.0 ps from the final 160 ns of the 200 ns trajectories. These trajectory-averaged residue-wise RMSF values were then used to calculate their mean and standard deviation across the five independent runs.

## Results and Discussion

To gain comprehensive insights into the dynamic structural changes of syntenin-1, we conducted a statistical comparison of the crystal structures obtained. As depicted in Figure 1, syntenin-1 molecules form dimeric assemblies within the crystal lattice, resulting in each asymmetric unit containing four PDZ domains (two PDZ1 and two PDZ2 domains from two chains). From a total of 10 determined crystal structures, we obtained 20 structures for the PDZ1 domain and 20 structures for the PDZ2 domain. The PDZ1 domain encompasses residues Glu114 to Arg193 (colored orange), while the PDZ2 domain spans residues Arg197 to Phe273 (colored blue). The apo-syntenin-1 structure was determined using a 163-amino acid construct, whereas the syntenin-1-compound complex structure utilized a 166-amino acid construct. Then, comparative analysis of these ten PDZ1-PDZ2 tandem structures revealed significant conformational heterogeneity exclusively within the PDZ1 domain of chain B. In contrast, the PDZ1 domain of chain A and both PDZ2 domains (chains A and B) exhibited structural conservation, as evidenced by superimposition analyses (Figures 2A, B, D, E). To quantify this observed divergence, we conducted statistical evaluations of root-mean-square deviation (RMSD) of backbone Ca positions. Initially, we classified these crystal structures based on their structural similarity. Pairwise RMSD values were computed for all possible pairs derived from the 20 PDZ1 as well as 20 PDZ2 domain structures. The resulting two-dimensional data matrix of pairwise RMSD was subjected to hierarchical clustering in both rows and columns. The data were sorted by closely related structural clusters in proximity. A heatmap was then generated, with color-coding indicating the level of pairwise RMSD (Figure 3).

**Figure 2:**
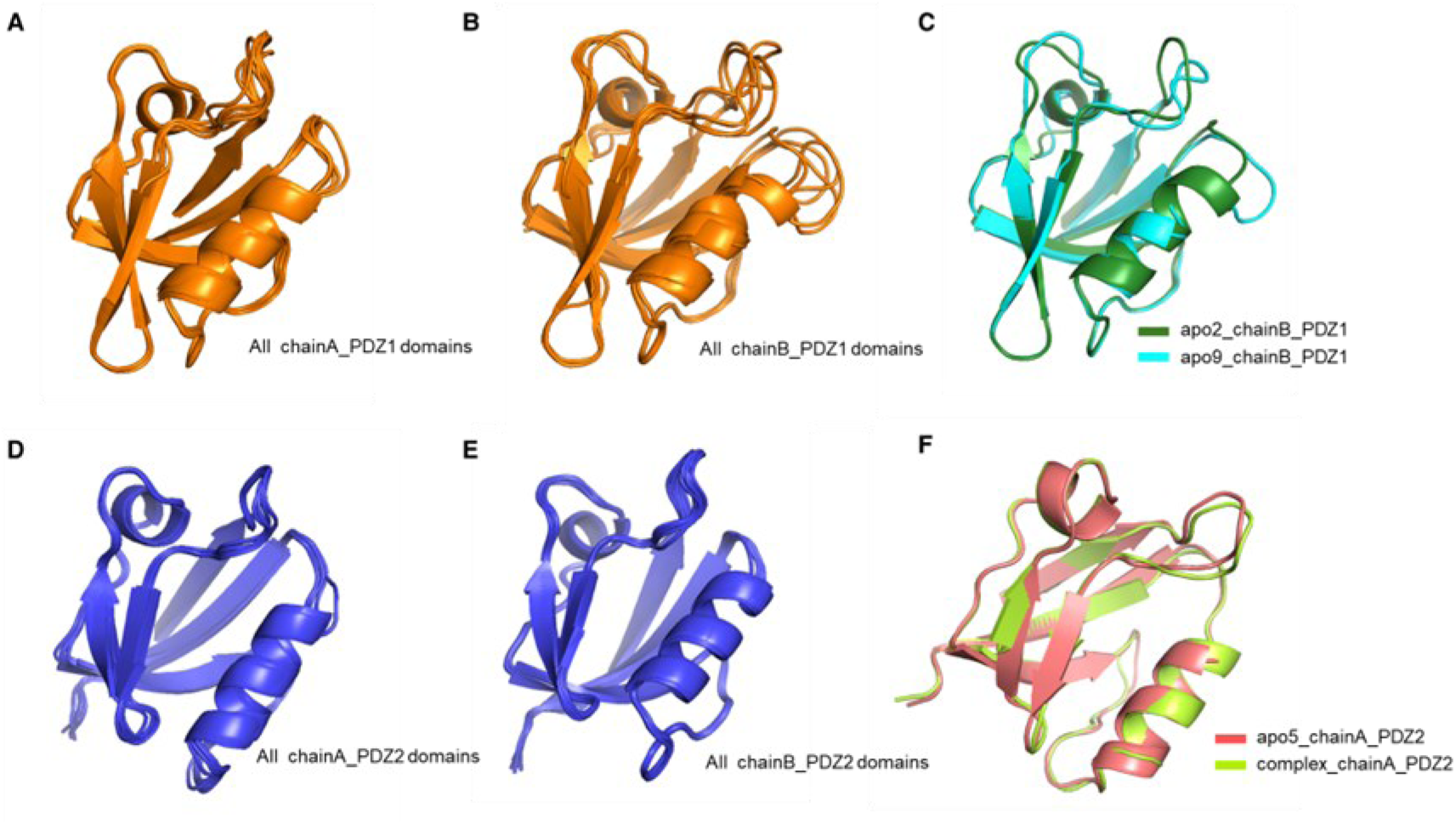
Comparison by superposition between each PDZ domain. A: Comparison of all nine structures of the apo PDZ1 domain of chain A. B: Comparison of all nine structures of the apo PDZ1 domain of chain B. C: Comparison of the pairs with highest pairwise RMSD among PDZ1 domain of chain B. D: Comparison of all nine structures of the apo PDZ2 domain of chain A. E: Comparison of all nine structures of the apo PDZ2 domain of chain B. F: Comparison of the bound and unbound structures of the PDZ2 domain.

**Figure 3:**
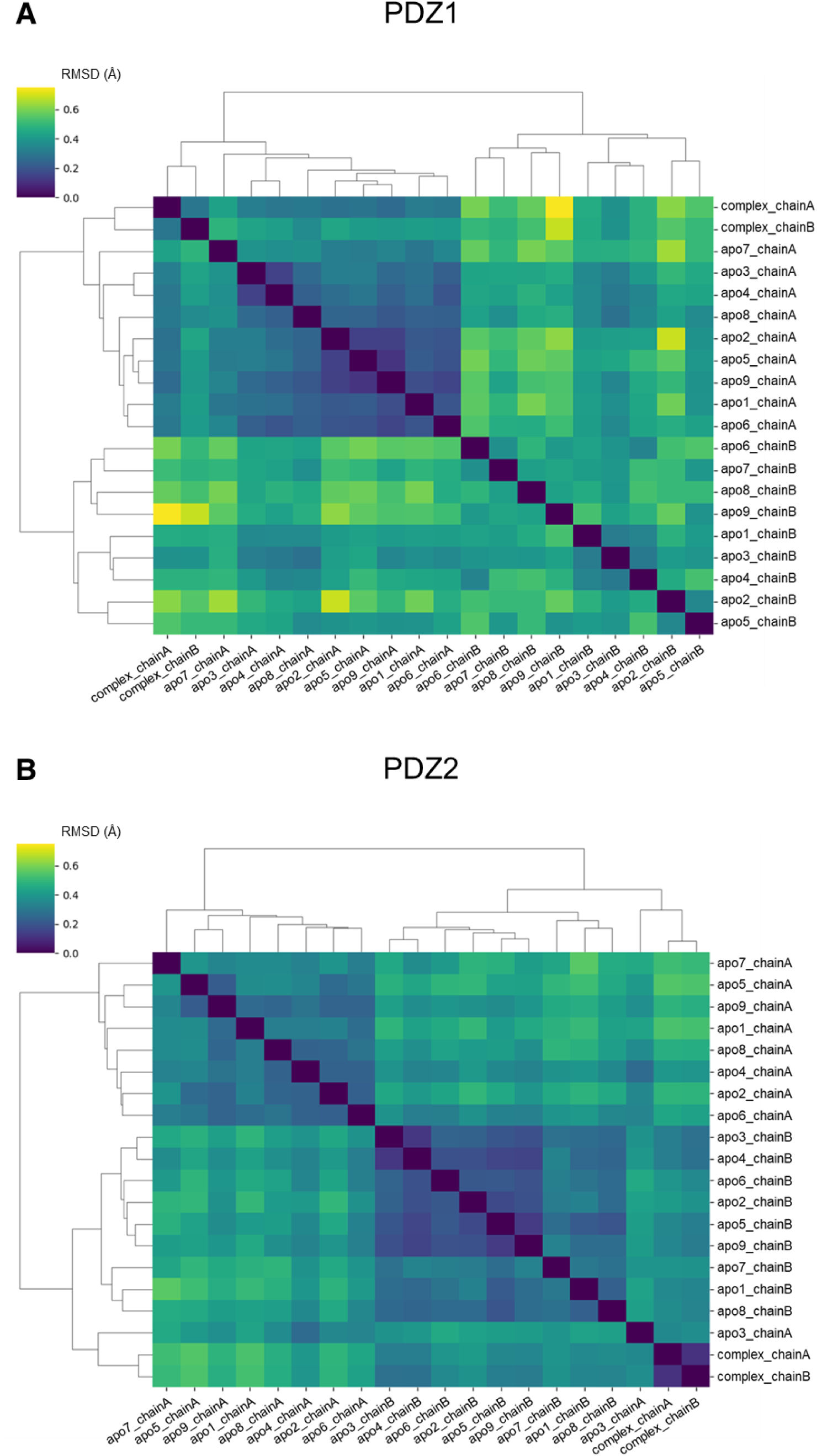
Cluster analysis of syntenin-1 PDZ domains. The structures of PDZ domains were extracted one by one from 9 apo structures and 1 complex structure, and the structures of 20 PDZ domains were obtained respectively. Pairwise RMSD was calculated for all pairs produced from 20 PDZ domains. Pairwise RMSD levels were classified from yellow to dark blue, and the similarity between each pair was represented in a heatmap. Based on the similarity, the cluster analysis was performed, and the results were expressed in a cluster dendrogram. A: Cluster analysis of PDZ1 domains. Major clusters were formed in chain A and chain B, respectively. B: Cluster analysis of PDZ2 domains. Major clusters were formed in apo chain A and apo chain B. A minor cluster was formed by apo3 chain A and complex structures.

For the PDZ1 domain, major clusters were observed to form between chain A/chain B pairs (Figure 3A). It should be noted that the dominant structural difference between chain A and chain B probably arises from crystal packing contacts. We concluded that the overall structural difference observed between PDZ1_chain_A and PDZ1_chain_B DO NOT reflect the intrinsically dynamic nature of PDZ1. In contrast, relatively large structural fluctuations were observed within PDZ1 domains of chain B/chain B pairs, suggesting that the intrinsic structural flexibility of the syntenin-1 PDZ1 domain becomes apparent in this context. Among this cluster, however, apo6 and apo7, which were crystallized without compounds, fell into the same structural subcluster, apart from the other subclusters that were crystallized with NPL inhibitors. Thus, we could not rule out that presence of inhibitors in crystallizing conditions exert some influence on structural differences of PDZ1. Conversely, PDZ1 of chain A/chain A pairs did not exhibit remarkable pairwise RMSD values, indicating generally similar structures across these instances (Figure 3A). Notably, the presence or absence of a compound in the crystallization condition did not influence the clustering of chain A/chain A pairs. This suggests that small structural diversity was primarily due to the restrictions imposed by crystal contacts and not affected by presence of absence of the PDZ inhibitors.

Next, we analyzed the structural differences of PDZ2 domains, which also exhibited two dominant clusters based on the difference between chain A and chain B (Figure 3B). However, unlike PDZ1, the hierarchical clustering analysis of PDZ2 structure succeeded in separating structures of apo3_chain A and both chain A and chain B of the complex structures into distinct major clusters. This result can be explained by the binding of the inhibitor PDZ2i, which may fix the loop conformation. However, interestingly, we observed that the apo3_chain A structure is very similar to that of the complex structures. Thus, we expected that “conformational selection” mechanism may work rather than “induced fit” upon PDZ2i binding to PDZ2 of syntehin-1. Accordingly, when comparing the structural differences among PDZ1 and PDZ2, PDZ1 consistently exhibited higher pairwise RMSD values than PDZ2. This observation strongly led us to hypothesize that PDZ1 possesses greater structural flexibility than PDZ2.

Accordingly, we visualized the selected pairs of PDZ domains and structural clusters by superimposing (Figure 2). The pair apo2_chainB_PDZ1/apo9_chainB_PDZ1 displayed the largest pairwise RMSD value among all PDZ1 pairs (Figure 2C). The PDZ1 domain demonstrated diverse structures in the loops surrounding the canonical ligand binding pocket. As mentioned previously, greater structural polymorphism was consistently observed in PDZ1 domains of chain B compared to those of chain A, when all obtained PDZ1 structures within chain A and chain B had compared each other, respectively (Figure 2A, 2B). In contrast, comparisons of PDZ2 domains within both chain A and chain B (Figure 2D, E) revealed no significant structural polymorphism, suggesting a rigid structure for PDZ2. Furthermore, a comparison between the bound and unbound structures of the PDZ2 domain (Figure 2F) exhibited only a small structural difference in a loop region analogous to where polymorphism was observed in the PDZ1 domain.

To analyze this structural diversity in detail, we calculated the RMSD for each residue within the syntenin-1 PDZ domains. For all PDZ domain pairs depicted in Figure 2, the pairwise RMSD for each residue was calculated, and their average values were plotted against the residue numbers (Figure 4). For PDZ1, locally high average pairwise RMSD values were observed in the Lys119-Ile125 loop and the Ala181-Glu184 loop, which are surrounding the canonical ligand binding pocket (Figure 4A). This suggests that these regions are intrinsically flexible. Additionally, minor peaks were observed at Ala143 and Leu149. In contrast, for PDZ2, although peaks were present in the aforementioned loops, they did not exhibit the same high average pairwise RMSD values observed for PDZ1 (Figure 4B). Small peaks were also observed at Asp224 and Gly243. Overall, the crystal structure analysis clearly indicated that PDZ1 exhibits higher structural flexibility than PDZ2, particularly in the identified loop regions.

**Figure 4:**
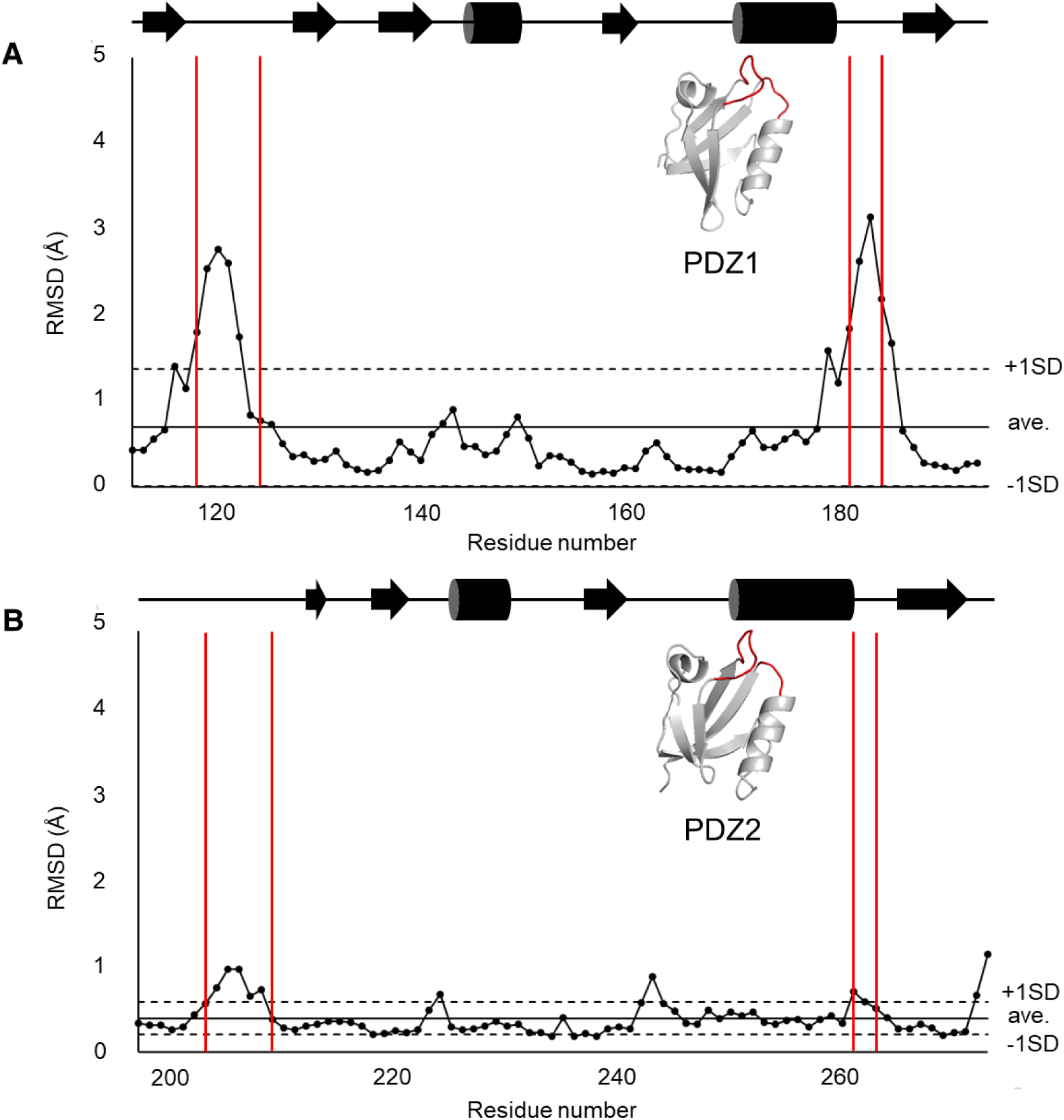
Average pairwise RMSD for each residue calculated by comparison of crystal structures. Highlighted areas (red) are the sites of molecular fluctuations in the PDZ1 domain. A: Average pairwise RMSD for each residue in the PDZ1 domain. B: Average pairwise RMSD for each residue in the PDZ2 domain. The diagrams of the secondary structures were shown in the top of the panels.

The structural heterogeneity observed in PDZ1 loop regions across ten crystallographic datasets prompted the hypothesis that these loops exhibit inherent conformational flexibility in solution, a property potentially linked to syntenin-1’s multifunctional role in cellular processes. To evaluate this hypothesis, we conducted molecular dynamics (MD) simulations of isolated human syntenin-1 PDZ domains using the GROMACS software package (Figure 5). Comparative analysis revealed enhanced flexibility in PDZ1 loop regions relative to PDZ2, consistent with crystallographic observations. Strikingly, simulations identified two previously uncharacterized regions of elevated mobility: (region i) residues Ile132-Gly135 in PDZ1 and (region ii) residues Asn230-Leu232 in PDZ2. Both regions are localized to the PDZ1-PDZ2 interdomain interface (Figure 6), with >80% of constituent residues participating in interdomain contacts. In detail, mainchain and side chain of Asp133 in region i of PDZ1 formed hydrogen bonds with Thr234 in PDZ2 (Figure 6A). Similarly, Gly231 in region ii in PDZ2 formed a hydrogen bond with Gln157 PDZ1 (Figure 6B). Thus, these regions may have additional intrinsic flexibility, although they adopted rather fixed conformations in tandem PDZ1-PDZ2 domain architecture of human syntenin-1.

**Figure 5:**
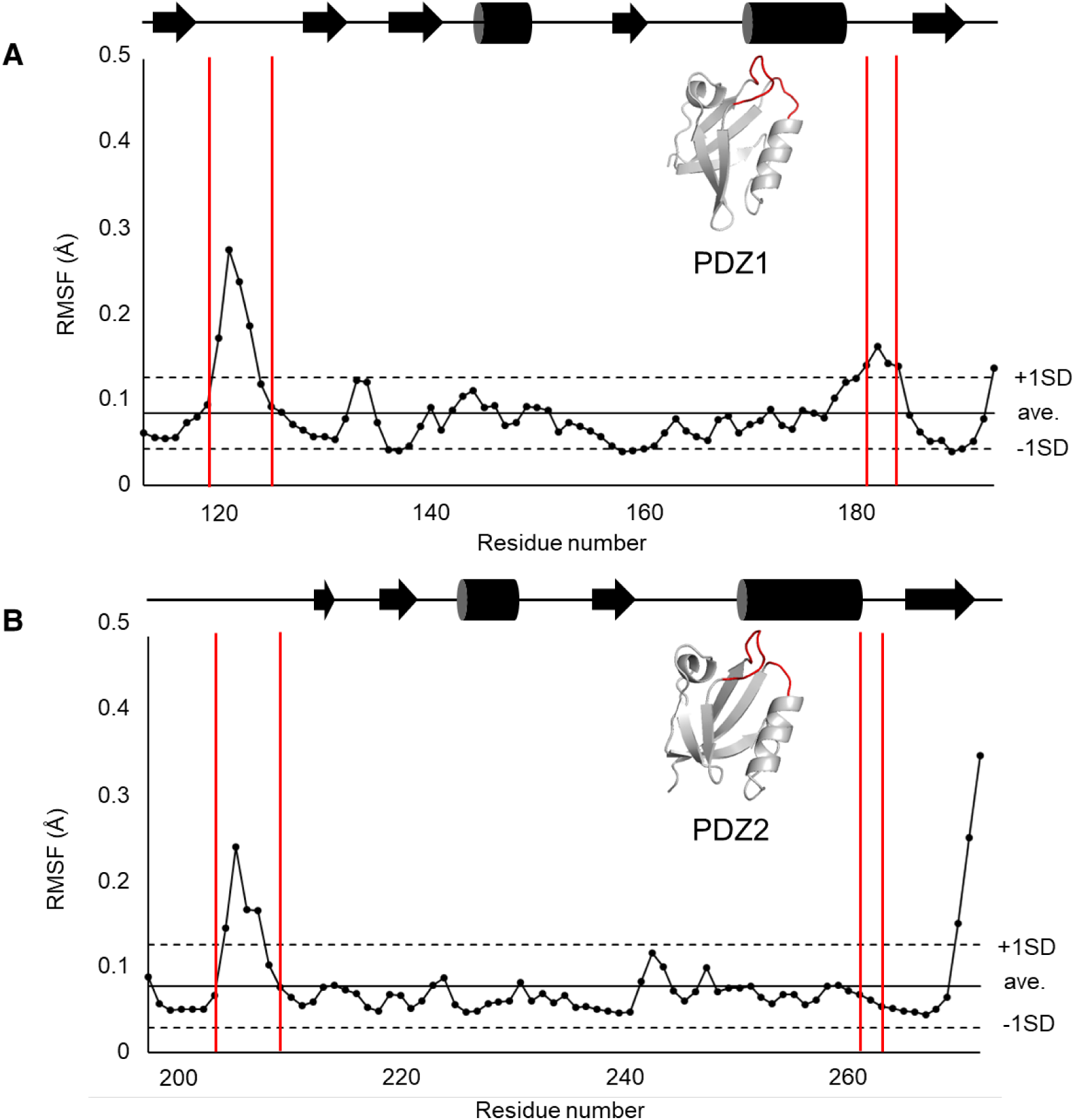
RMSF for each residue calculated by MD calculations. Highlighted areas show molecular fluctuations in the PDZ1 domain. A: RMSF for each residue in the PDZ1 domain. B: RMSF for each residue in the PDZ2 domain. The diagrams of the secondary structures were shown in the top of the panels.

**Figure 6:**
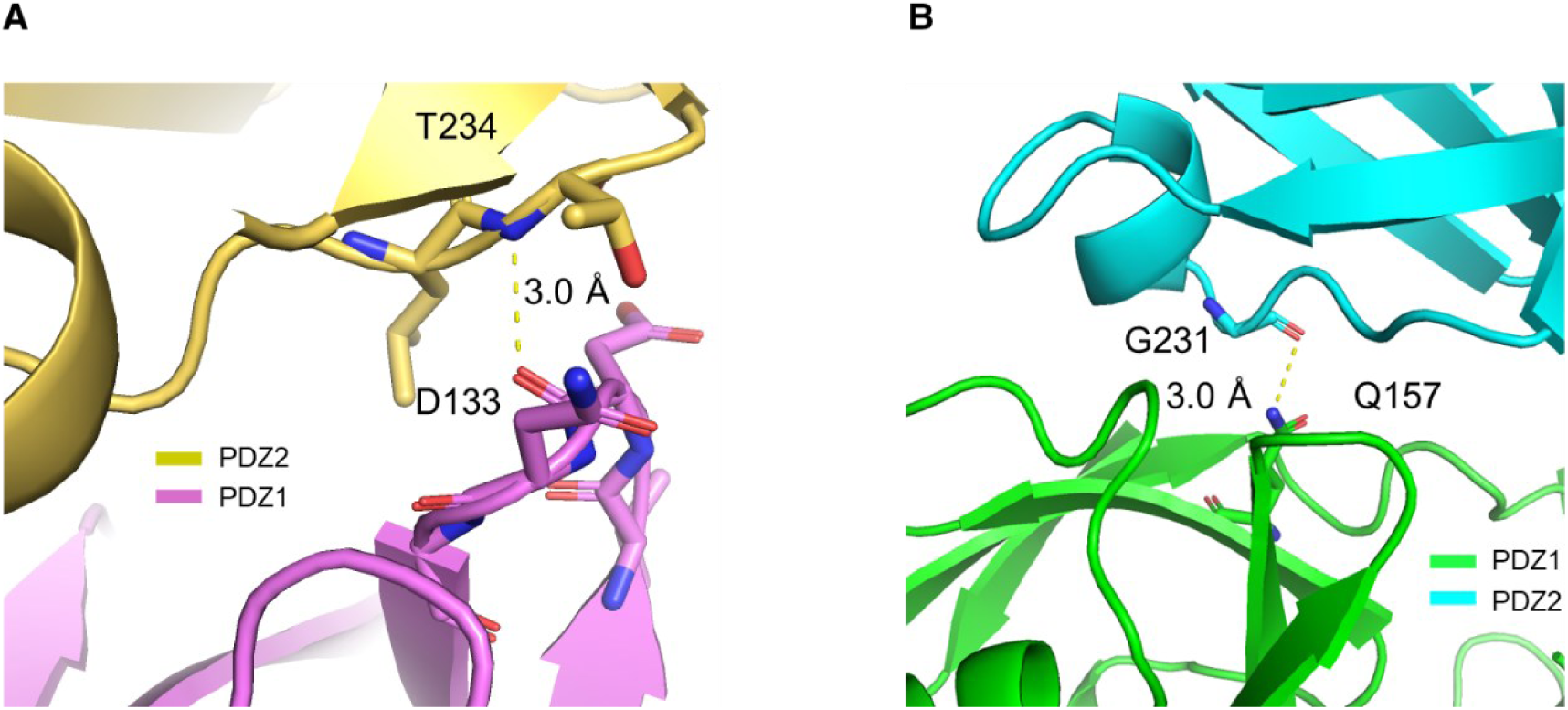
The residues that exhibited a high RMSF in MD simulations with a low structural diversity in the ten crystal structures. A: Around Ile132 to Gly135 in PDZ1. B: Around Asn230 to Leu232 in PDZ2.

## Conclusion

This study provides compelling evidence that PDZ1 in human Syntenin-1 exhibits inherent conformational diversity compared to PDZ2’s rigid architecture, despite their sequential similarity. Through extensive crystallographic analysis and molecular dynamics simulations, we demonstrate that PDZ1’s structural plasticity is concentrated in key ligand-binding loops, suggesting a conformational selection mechanism for diverse partner recognition. In contrast, PDZ2 maintains remarkable structural conservation, indicating distinct evolutionary constraints on these tandem domains. The observed dynamic asymmetry explains Syntenin-1’s versatility in cellular signaling pathways and offers a rational foundation for developing domain-selective inhibitors targeting cancer metastasis, viral pathogenesis, and neurodevelopmental disorders. Our findings highlight the importance of multi-structure comparisons and hierarchical analysis for accurately mapping protein dynamic landscapes, beyond what B-factors alone can reveal.

## Author Contributions

Conceptualization, T.T., N.I., and H.H.; Investigation, N.A., Y.H., K.S., N.N. and T.T.; Graphics, Table and Charts, N.A. and K.S.; Data Analysis, N.A., Y.H., K.S., N.N. and T.T.; Resources, Y.H., K.S., N.N., T.T.; Writing and Original Draft Preparation, N.A. and H.H.; Writing, Reviewing and Editing, N.I. and H.H.; Supervision, N.I. and H.H.; Project Administration, H.H.; Funding Acquisition, H.H. All authors have read and agreed to the published version of the manuscript.

## Acknowledgments

The synchrotron radiation experiments were performed with the approval of the Photon Factory Program Advisory Committee (Proposal No. 2022G024, No. 2024G043).

## Data Availability Statement

The datasets of crystallographic analyses are deposited to the Protein Data Bank. Other related data (MD trajectories) will be available at the request to authors.

