## Supplementary Figure S1 for "Intrinsically Dominant Conformational Diversity in PDZ1 within the Tandem PDZ1-PDZ2 of Human Syntenin-1 Underlined by Crystal Structures"

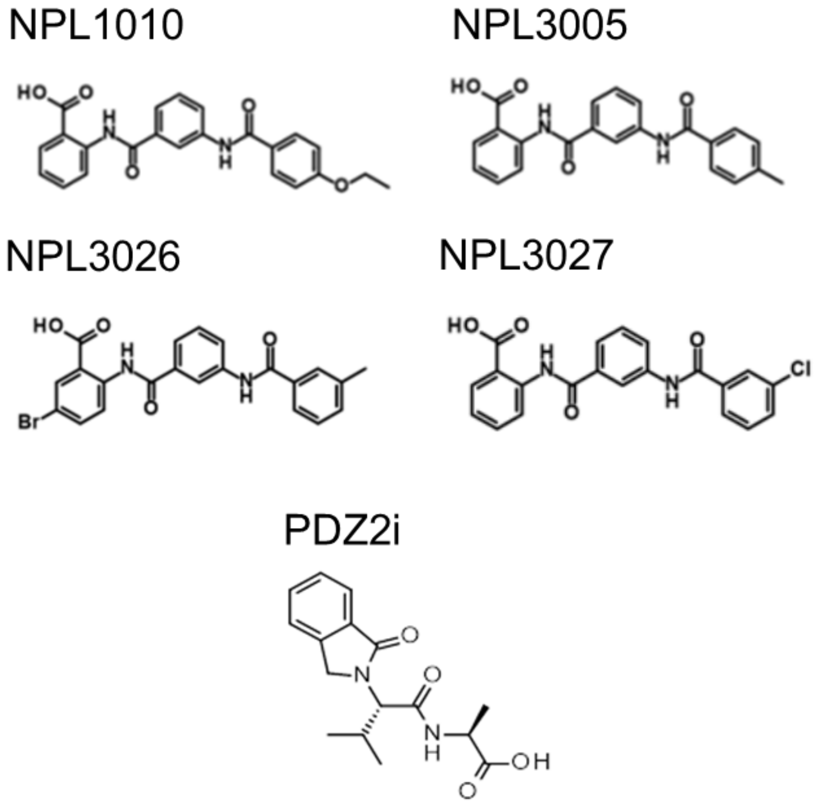


**Supplementary Figure S1.**

The chemical structures of the compounds used in crystallization experiments of human syntenin-1 PDZ1-PDZ2 tandem.
